# Theta and gamma oscillations in the rat hippocampus during attentive lever pressing

**DOI:** 10.1101/243527

**Authors:** A Lipponen, H Tanila, K Gurevicius

## Abstract

The hippocampus is known to be pivotal for spatial memory but emerging evidence suggests its contribution to temporal memories as well. However, it is not clear how the hippocampus represents time and how it synchronizes spatial and temporal presentations into a coherent memory. We assessed the specific role of hippocampal theta and gamma oscillations and their interaction in short-term timing of motor reactions. Rats were trained to maintain lever pressing for 2.5 s and then to quickly release the lever and retrieve water reward from a nearby water port guided by a cue light. In essence, this task allows observation of hippocampal rhythms during timed anticipation when no overt movements take place. Then we implanted wire electrodes to five hippocampal layers for recording local field potentials during the task. Consistent with earlier reports, theta showed a declining trend during the lever press. We also found that fast-gamma declined in tandem with theta while slow-gamma showed an opposite trend. Theta-phase to gamma-amplitude cross-frequency coupling measured with modulation index (MI) varied significantly between the three task phases. Interestingly, also changes in MI were opposite for fast- and slow-gamma. The MI was also related to the task performance, so that during omission trials the MI for fast-gamma in CA1 was smaller than during trials with premature lever release. In addition, the MI in dentate hilus was higher during all error trials than during correctly performed trials. Collectively, these data suggest an important role of synchronization of hippocampal theta and gamma rhythms to timing of cued motor reactions.

## Introduction

Studies in human patients with medial temporal lobe damage as well as lesion studies in experimental animals have established the pivotal role of the hippocampus in memory functions, especially in long-term declarative memory (Scoville and Milner, 1957; Morris et al., 1982; Corkin, 1984; Squire, 1992; Corkin, 2002). On the other hand, functional measurements of hippocampal activity in rodents revealed hippocampal neurons to respond only when the animal is situated in a particular area of the testing platform, leading to the idea that the main function of hippocampus and connected areas is to store information of spatial context, thus providing framework for a cognitive map (O’Keefe and Dostrovsky, 1971; O’Keefe and Nadel, 1978; Fyhn et al., 2004).

Therefore, it has been a challenge to understand how hippocampus is able to establish not only spatial but also temporal frameworks for episodic memory. It is assumed that a single mechanism involved in spatial memory serves also temporal framework. According to this hypothesis a subset of neurons that fire during given gamma cycle, i.e. cell assembly, form a pattern representing an item (episodic memory) similarly as spatial location (Buzsáki, 2005; Buzsaki and Moser, 2013). Sequential activation of these assemblies is replayed repeatedly within single theta cycles with the same assembly occupying different phases of the cycle. This mechanism could thus provide the crucial binding of individual details into a coherent memory of an event (Buzsáki, 2005; Buzsaki and Moser, 2013). According to another hypothesis, place cell activity is also associated with time (MacDonald et al., 2011; MacDonald et al., 2013). As such these so-called time cells encode moments in a temporally structured experience as spatial cells code for locations, and the activity of these two cell types thus reflects only different dimensions of the context in which learning occurs (MacDonald et al., 2011; MacDonald et al., 2013).

Based on these findings we hypothesized that temporal aspect of hippocampal functions might be related not only to the activity of individual neurons but also to the rhythmic activity of hippocampal neuronal ensembles, i.e. brain rhythms (Buzsaki et al., 2012). The two most prominent hippocampal brain oscillations are the theta (4–12Hz) (Green and Arduini, 1954; Weiss and Fifkova, 1960) and gamma rhythms (25–90 Hz) (Stumpf, 1965; Leung et al., 1982). Even though these rhythms have been strongly linked to several kinds of behavior (Vanderwolf, 1969; Whishaw and Vanderwolf, 1973; Kramis et al., 1975) their exact link to mnemonic functions of the hippocampus is still arguable. For instance, lesioning of the medial septum abolishes hippocampal theta and impairs learning and recalling of locations in a spatial memory task, but if lesioning is performed before training it does not totally prevent spatial learning (Winson, 1978). However, it is plausible that the coherent interactions of hippocampal rhythms are well suited for the regulation of the spatial and temporal coordination of information flow and processing (Canolty et al., 2006; Canolty and Knight, 2010). For example, the theta phase is known to modulate gamma power in rodent hippocampal and cortical circuits (Buzsaki et al., 1983; Bragin et al., 1995; Jensen and Colgin, 2007), and this theta-gamma coupling has been shown to have a functional role in the memory functions of hippocampus (Tort et al., 2008; Shirvalkar et al., 2010).

The present study delved into the timing function of the hippocampus. We first developed a simple lever pressing task in which animals were required to press a lever for at least 2.5 s to receive a reward. The task design was to minimize the confounding effect of exploration and associated movement-related theta and to focus on the timing function (Vanderwolf, 1969; Whishaw and Vanderwolf, 1973; MacDonald et al., 2013). Secondly, we tested the hypothesis that successful encoding of temporal requirements of this task can be predicted by changes in hippocampal theta and gamma rhythms. However, as it has been established that these oscillatory activities are not independent, we also assessed modulation of gamma oscillations by the theta rhythm (Tort et al., 2008; Tort et al., 2010).

## Methods

### Animals

The subjects were male Wistar rats (n=10, weight 324 ± 0.4 g, mean ± SEM). The rats were caged after operation individually in a controlled environment (temperature +21± 1 ° C, light on 7:00 – 19:00) with water and food available *ad libitum*. During the behavioral training and recordings water was removed before each session for 8–12 hours to increase motivation. All experiments were conducted in accordance with the guidelines of the Council of Europe and approved by the State Provincial Office of Eastern Finland.

### Electrode implantation

The surgery was performed under isoflurane gas anesthesia (airflow 450 ml/min, 4.5 % for induction and 250 ml/min, 2.1 % for maintenance). The rats were implanted with wire electrodes (Formwar^®^ insulated stainless steel wire, diameter 50 μm, California Fine Wire Company Co, Grover Beach, CA, USA with vertical tip separation of 600 μm) in the following hippocampal coordinates: CA1: AP -3.6 mm (from bregma), ML 1.6 mm (from the midline), DV 2.8 mm (from dura); CA3: AP -3.6, ML 3.6, DV 3.6; and DG: AP -4.2, ML -1.4, DV 4.2). A parietal screw served as a ground and a frontal screw as a reference. After the surgery, the rats received carprofen (5mg/kg, i.p., Rimadyl^®^, Vericore, Dundee, UK) for postoperative pain alleviation. During the following 2–3 days the animals were given carprofen 5 mg/kg/day in drinking water, and antibiotic powder (bacitrasin 250 IU/ g and neomycinsulfate 5mg/g, Bacibact^®^, Orion Finland) was applied, if necessary, onto the wound.

### Behavioral task

After a two-week recovery period the rats were trained to perform the lever pressing task for water reward during 20–30 sessions. In this task that was designed to assess the plausible timing function of hippocampus, the animals needed to press a lever uninterruptedly for 2.5 s to activate a water delivery system. The task took place in a plastic cage (width 31.5 cm, length 34 cm and height 40 cm) equipped with a metal lever (length 3.4 cm, height from the floor 2 cm) and a water port on the same side of the cage. The lever and the water port were separated by a transparent glass wall protruding 10 cm from the wall forcing the rat to run to the water port after releasing the lever instead of just turning the upper body. The task was guided by two signal lights, one cue light above the lever and a second light over the water port. The task started with the cue light turning on. After a constant 2.5 s of lever pressing the cue light was switched off while the light above the water port was switched on providing an external cue for the animal indicating successful trial (Fig. 1.). After a nose poke to the water port, the light above the port was switched off while the cue light was switched on to indicate the possibility of a new trial.

**Fig. 1.**
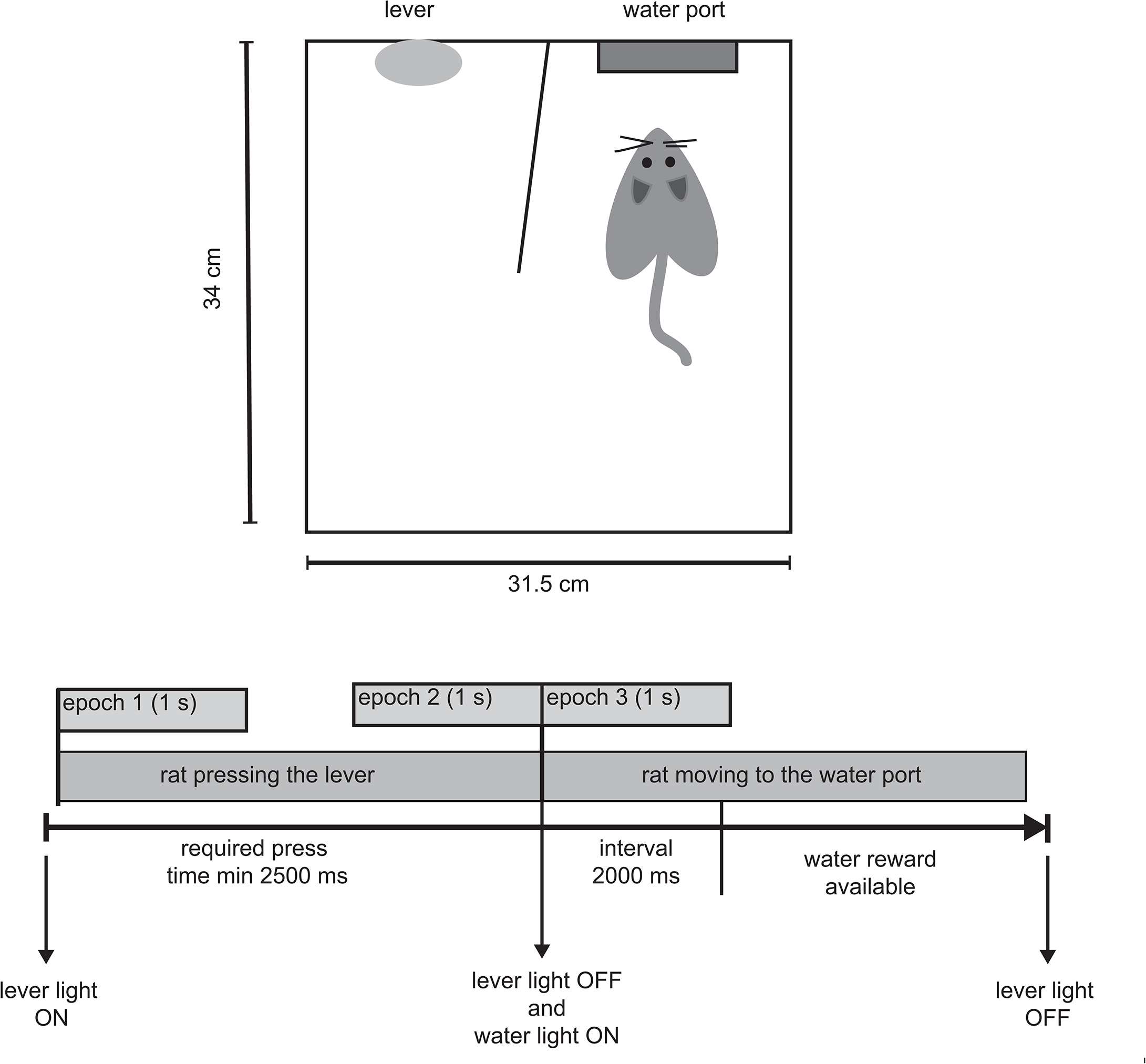

The training consisted of four different phases. In Phase 1, the rat was accustomed to the operant box. In addition, the light above the water port was switched on for 2 s to attract the attention of the animal, and the water drops were delivered to the water port. The water delivery took place at randomized 10 to 30-s intervals. The lever was fixed in this phase of training. In Phase 2, the rat was accustomed to lever pressing. Now the light above the lever was constantly on and the water drops were delivered between random intervals directly on the lever. Occasional visits to the water port had no consequences. The training finished after ~80 drops were delivered during a 30 to 40 min training session. In Phase 3, the rat needed to activate the water port by pressing the lever for at least 200 ms. In Phase 4, duration of lever pressing was gradually increased from 200 ms in 200-ms steps after every 3 repeated successful trials. When the animal reached a stable 1800 – 2000 ms lever pressing time, a fixed pressing time of 2500 ms was introduce. The rat was repeating this final training stage as long as it reached a 60 % performance level (60 right performances in 100 trials) in four sessions in a row. Overall the training took 20–30 days.

### EEG recording

EEG was recorded during the task performance once the rat had attained the criterion level performance. The signal from the recording electrodes passed through a unity-gain preamplifier (Neuralynx Inc., Bozeman, MT, USA) mounted on the distal end of the recording cable to remove movement artifacts and further through the main amplifier (gain 1000, band-pass filtering 1 Hz – 1 kHz, A-M systems Inc., Sequim, WA, USA). The signal was digitized and acquired with the SciWorks system (DataWave Technologies, Longmount, CO, USA). A + 5 V analog signal was directly fed into one data channel when the lever was pressed and closed a separate battery-powered circuit. The timing of light signals and water delivery was controlled by another desktop computer with a custom-made program.

### Histology and verification of electrode locations

After the experiments the animal received an overdose of medetomidine (1 mg/ml) and ketamine (50 mg/ml) mixture (0.5 ml/kg, 0.8 ml med. + 2.4 ml ket., i.p.). The locations of electrodes were marked by passing 50 μA DC current 8–10 s through the electrodes. Then the animal was transcardially perfused firstly with cold saline for 5 min at 13 ml/min followed by 10 % formalin solution for 13 min at 13 ml/min. The brain was moved from the skull and left for immersion postfixation in 10% formalin for 4 h, followed by 30 % sucrose solution for 2 days. The brains were stored in -20 ° C until slicing. A serial of coronal sections (thickness 35 μm) were cut with freezing slide microtome. The electrode locations were verified from the slices by cresyl violet – Prussian blue staining.

In addition to histological marks, we used well-known hippocampal electrophysiological markers to verify electrode location: theta and gamma power profile in hippocampus, phase of theta (Bragin et al., 1995), and phase of theta for ripples (Isomura et al., 2006). Electrodes with poor signal quality (due to bad contacts and electrical shortcut) or lack of theta-gamma modulation were excluded from further analysis.

### EEG analysis

Data from several sessions were pooled for the final analysis. All signals were normalized to amplification, and offline analyses were conducted using MATLAB (Mathworks, Natick, MA, USA). The power spectral density (PSD) for each epoch was estimated by using Welch’s averaged modified periodogram method of spectral estimation (0.5 s segments with 75% overlap) and averaged PSD per animal was calculated for each performance and each epoch. To eliminate sweeps with artifacts, we first calculated the distribution for averaged PSD values between 80 and 90 Hz. Then, outlying sweeps were excluded using iterative implementation of the Grubbs test for outliers (MATLAB routine “deleteoutliers.m” by Brett Shoelson). This effectively removed occasional artifacts related to bad contact, jerky movements, or animal handling.

The theta phase for gamma events was computed from event-related average triggered by event peaks (Bragin et al., 1995; Kramer et al., 2008). To quantify the gamma amplitude modulation by the theta phase, we estimated a modulation index (MI) based on a normalized entropy measure, as reported previously (Tort et al., 2008; Tort et al., 2010). This index is able to detect cross-frequency coupling between two frequency ranges of interest. We estimated the mean power of gamma oscillation for each 10 degree phase. Statistical significance of MI was estimated by creating shuffled versions of the time series (phase of one epoch and amplitude of another) and generated 200 surrogate MI values, where each MI value is calculated from 20 random pairs. Assuming a normal distribution of the surrogate MI values, a significance threshold was then calculated by using P < 0.01 as the threshold.

## Results

### Electrode locations

Electrode locations were verified by electrophysiological markers (e.g. CA1 ripple activity) and the post-mortem histological analysis, and were grouped into 6 distinct locations. CA1upper (n= 7) electrodes were located in the oriens or pyramidal cell layer of hippocampal CA1 area, CA1lower (n = 5) in stratum radiatum, CA1lm (n = 7) in lacunosum moleculare, CA3 (n = 5) in the pyramidal cell layer and stratum radiatum of CA3 area, PoDG (n = 8) in polymorph layer and MoDG (n = 9) in the molecular layer of the dentate gyrus (Fig. 2).

**Fig. 2.**
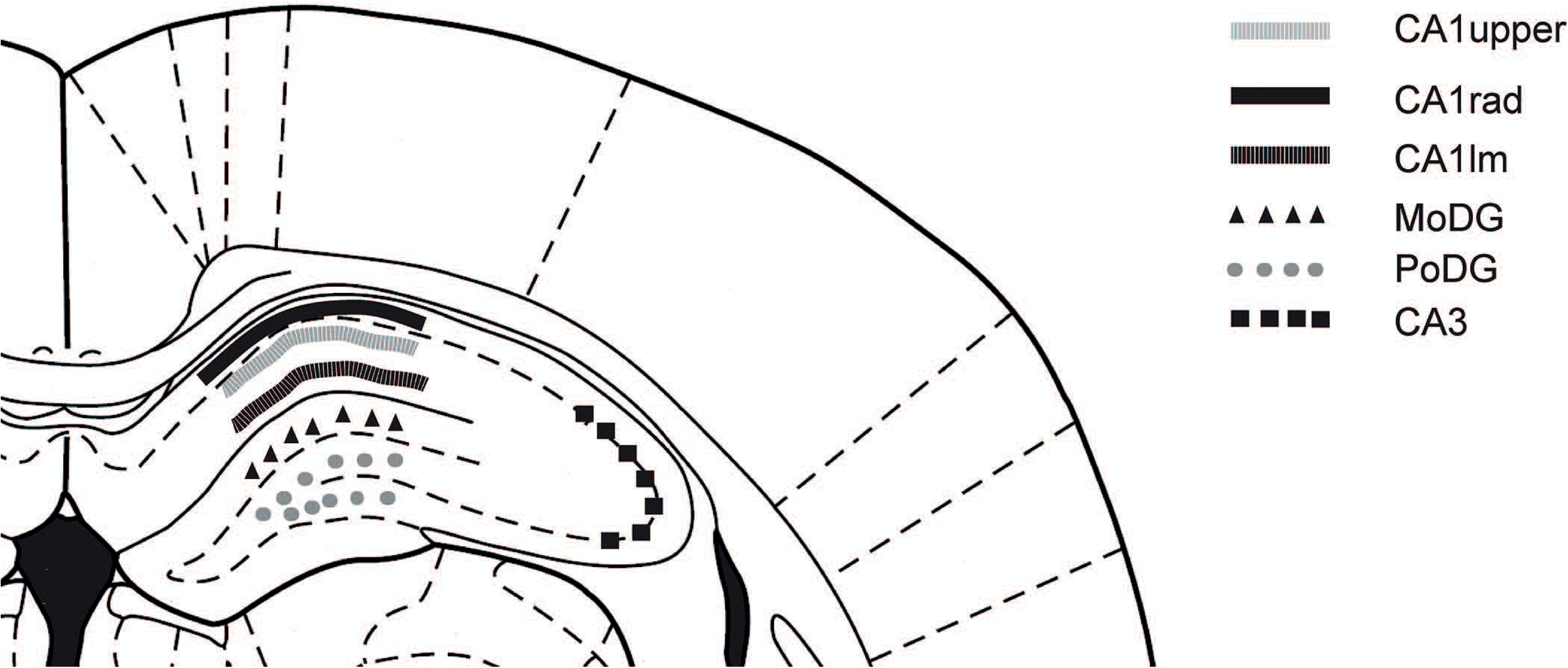

### Behavioral analysis and categories of trials and task epochs

Ten rats performed the task above the set criterion for the minimum of 5 consecutive days and had functional electrodes in the planned locations. All the animals accepted for further analysis reached the learning criteria of 60 % correctly timed responses during 4–5 consecutive sessions (Fig. 3).

**Fig. 3.**
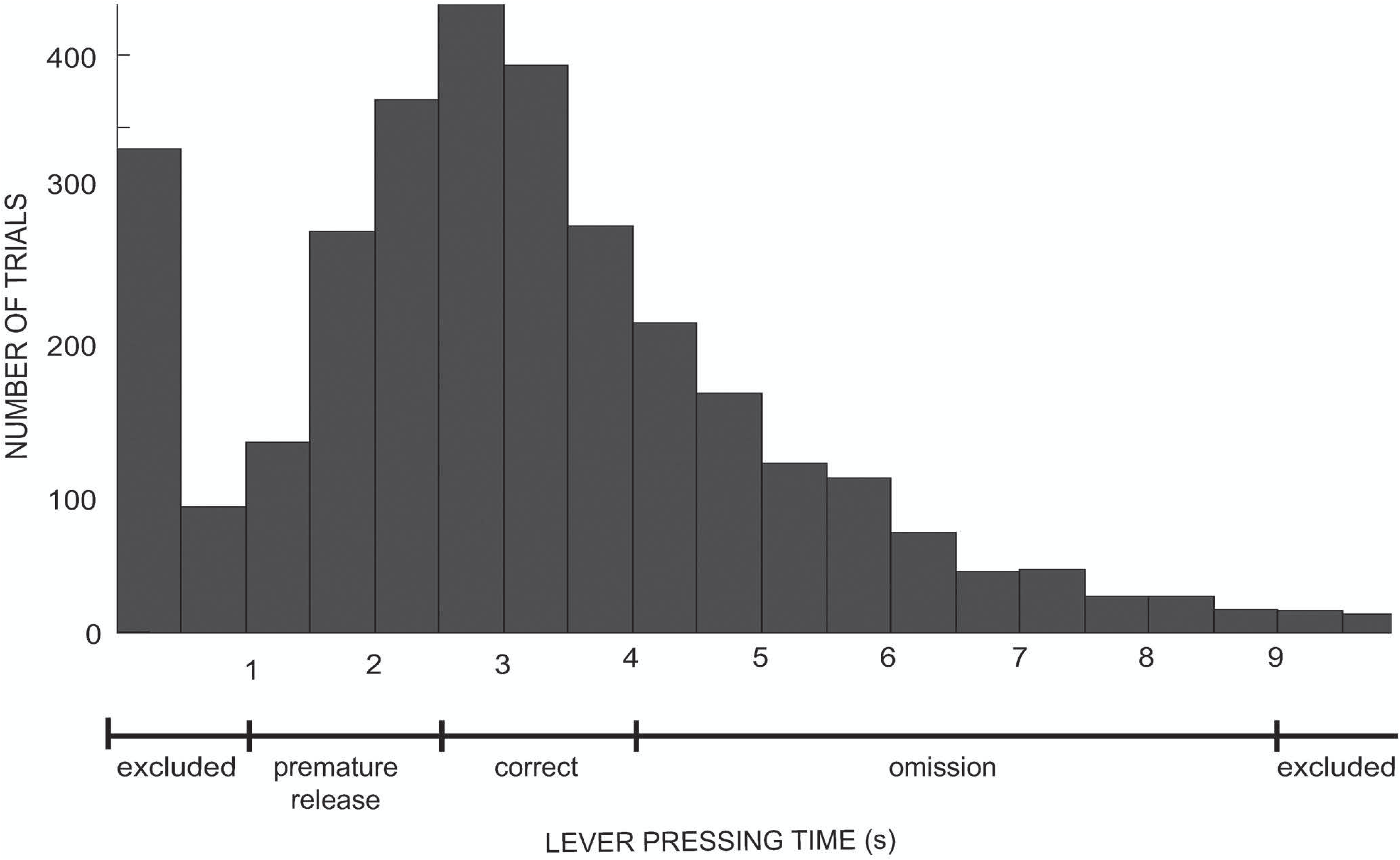

The rats performed two types of timing errors. The premature release trials consist of trials when the animal was pressing the lever for less than 2.5 s and, therefore, did not receive the water reward. However, in occasional trials the rat quickly pressed and released the lever in less than 0.5 s and then tried to immediately compensate this slip with longer pressing. Therefore, we excluded from the analysis trials when constant pressing was less than 1 s in duration, and, thus the premature category comprised pressing times from 1 s to < 2.5 s. Occasionally the rat simply remained lying on the lever for several seconds. We interpret these trials as non-attentive omissions with lever-press times between 4–9 s. All trials in which pressing time exceeded 9 s were excluded from analysis. Correct performance was thus strictly limited to lever-press times between 2.5 – 4.0 s.

The number of analyzed trials based on behavioral categorization were correct (n = 1113), premature release (n = 802) and omissions (n = 812). Each trial was further divided into 1-s epochs. Epoch 1 covered the first 1 s of lever pressing while Epoch 2 comprised the last 1 s before the lever release. Epoch 3 corresponds to the first 1 s after the lever release (see Fig. 1).

### Theta-oscillation shows a strong association with movement

Figure 4 illustrates that theta (6–10 Hz) oscillation was prominent in all hippocampal areas during all behavioral epochs (Fig. 4). Furthermore, theta oscillation fluctuated between the epochs, and similar changes were detected in all channels. The most prominent theta was present after the lever release (Epoch 3). This coincides with the movements of release and turning around. We observed stronger theta in the MoDG, PoDG and CA1rad channels during release epoch (Epoch 3) compared to the beginning of lever pressing (Epoch 1) (see table 1). In addition, during lever pressing (Epoch 1–2) theta power in CA1lm was significantly different during correct vs. error trials. In particular, the highest theta peak power was observed during premature trials while the theta power was smallest during omission trials (mean power at 7 Hz: correct = 1149 ± 368, premature = 1251 ± 395, omission = 1033 ± 323 mV^2^ *10^−6^) (Fig. 4 and Table 1). Similar results were observed also in CA1upper and CA1rad locations.

**Fig. 4.**
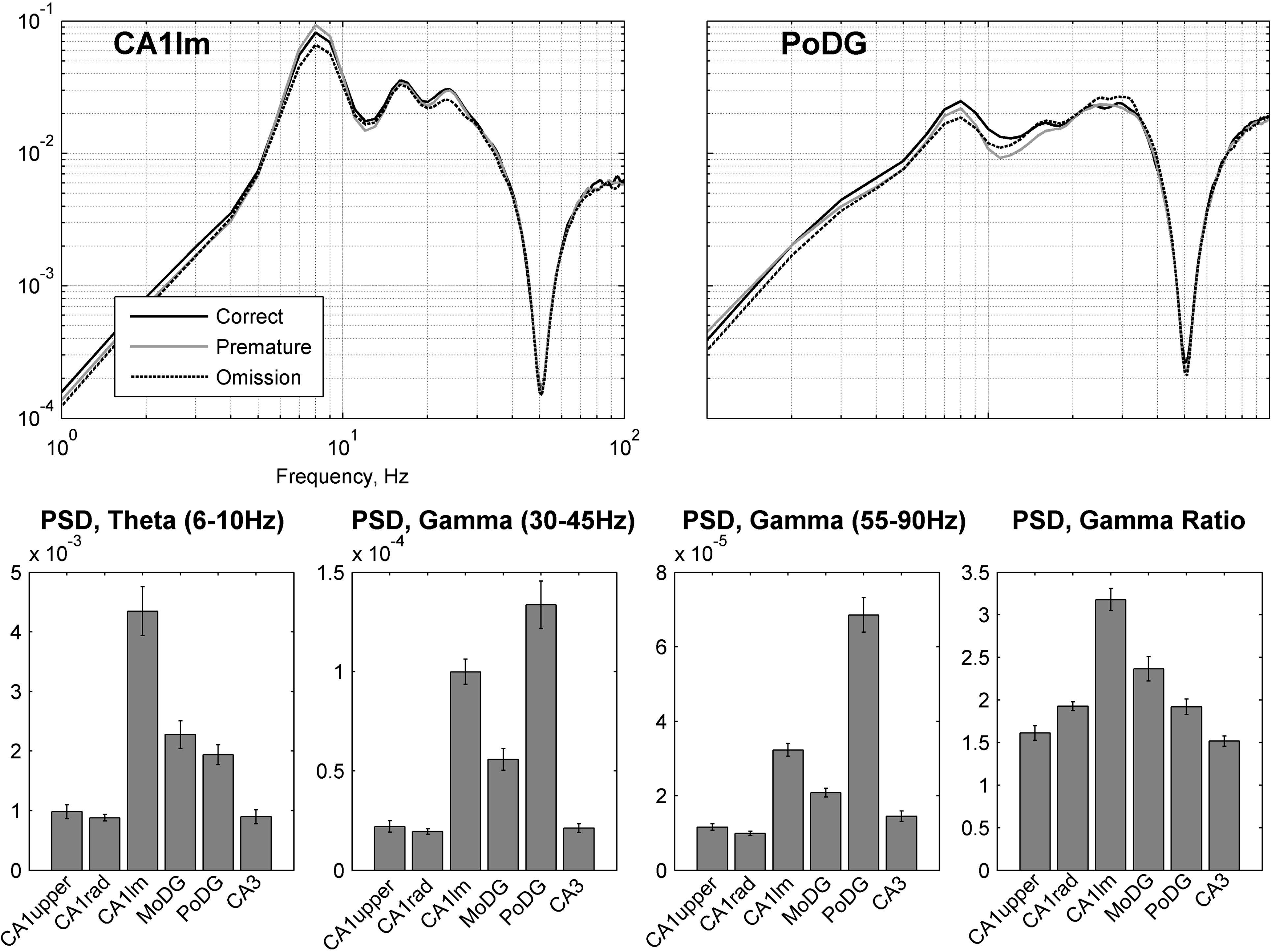

### Dynamic changes in gamma frequency range

We also observed task-related changes in the gamma frequency range. However, those changes were more location specific. Interestingly, task-related changes in gamma power were different for the slow (30–45 Hz) and fast (55–90 Hz) gamma range. Similar to theta oscillations the fast-gamma was increasing during lever pressing and releasing epochs as compare to Epoch1 (Fig. 5 bottom and table 1). In contrast task-related changes in the slow gamma power were opposite in direction to theta. That is, slow gamma power was strongest during Epoch 1 and smallest during Epoch 3 (Fig 5 top). Contrary to theta, there was no clear association between gamma power and performance accuracy except for slow-gamma in CA1lm during epoch 2&3 (Fig. 6 bottom and Table 1).

**Fig. 5.**
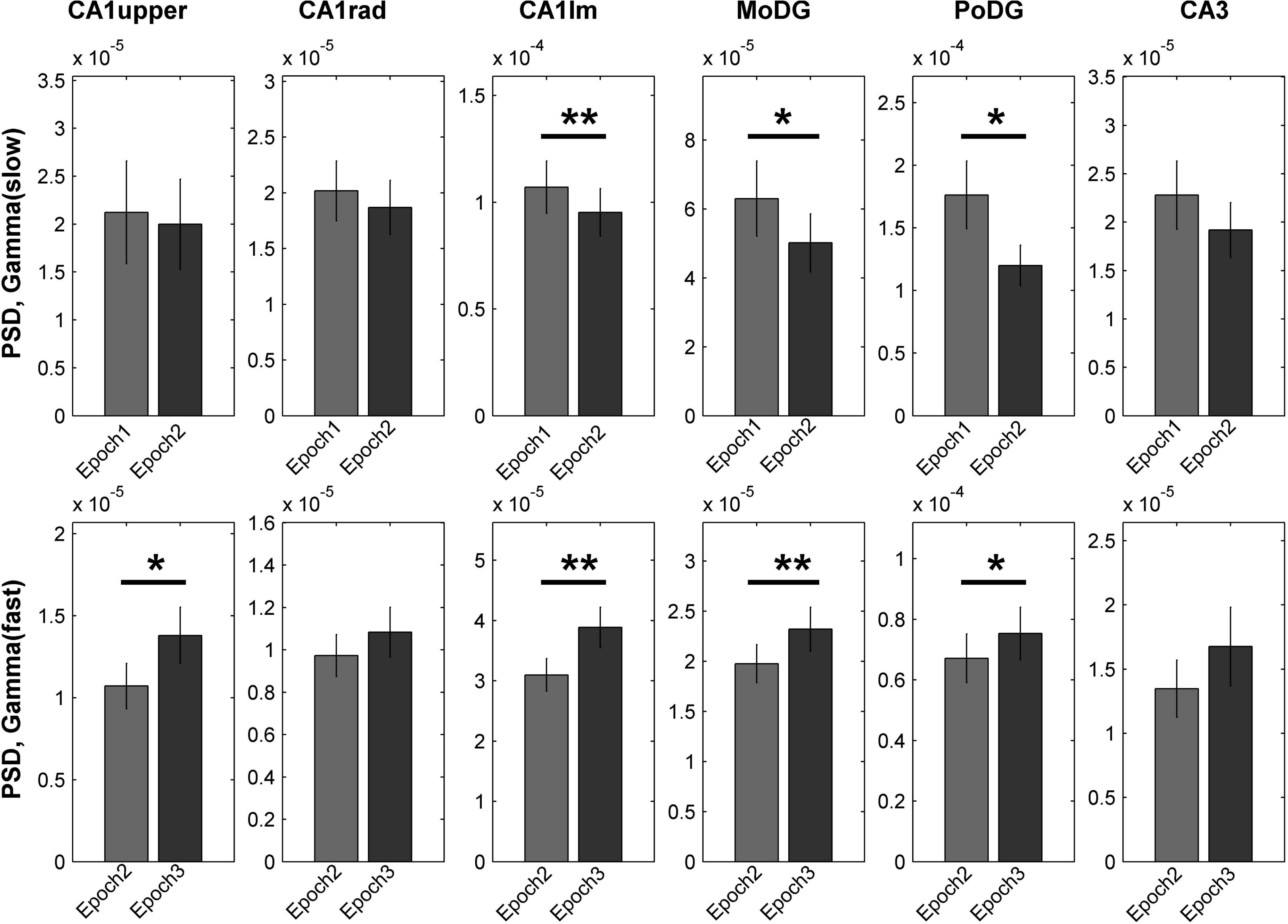

We further explored how slow- and fast-gamma interact in different hippocampal subfields by calculating the ratio between those two PSD values. The gamma ratio varied between hippocampal subfields and was highest in CA1lm (Fig. 4 bottom right). The gamma ratio showed task-related changes as well. It was significantly reduced in Epoch 2 when compared to Epoch 1, and further reduced during Epoch 3 (Table 2). This alternation in the gamma ratio was obvious in all recorded locations except CA1upper. The performance accuracy also affected the gamma-ratio. In particular in CA1lm, during lever pressing (Epochs 1&2) the gamma ratio was highest during premature trials and smallest during omission trials (Post**-**hoc p < 0.05, Table 2). Similarly, during lever release (Epochs 2 vs. 3) the gamma ratio in CA1lm and MoDG was higher during premature trials than during correct or omission trials (Post-hoc p < 0.05, Table 2). We also observed significant reduction of the gamma ratio during omission trials in PoDG during lever release (Epoch 2 vs. 3) (Post-hoc p < 0.05, Table 2).

### Task and performance specific alternation in cross-frequency coupling

We further tested the hypotheses that theta phase modulation of gamma amplitude in the two ranges (30–45 Hz and 55–90 Hz) (TG-Phase) differs between the task epochs and relates to performance accuracy. To test how gamma oscillations interact with theta, we plotted the gamma amplitude against the theta phase and measured a modulation index (MI) as described previously (Tort et al., 2008; Tort et al., 2010). First, we found that MI was above chance level (threshold of significance was set to p = 0.01) in all measured hippocampal locations (representative examples in Fig. 7). In addition, we observed specific alternations in the cross-frequency coupling related to epochs, and confirmed that these changes varied at the different recording locations.

**Fig. 6.**
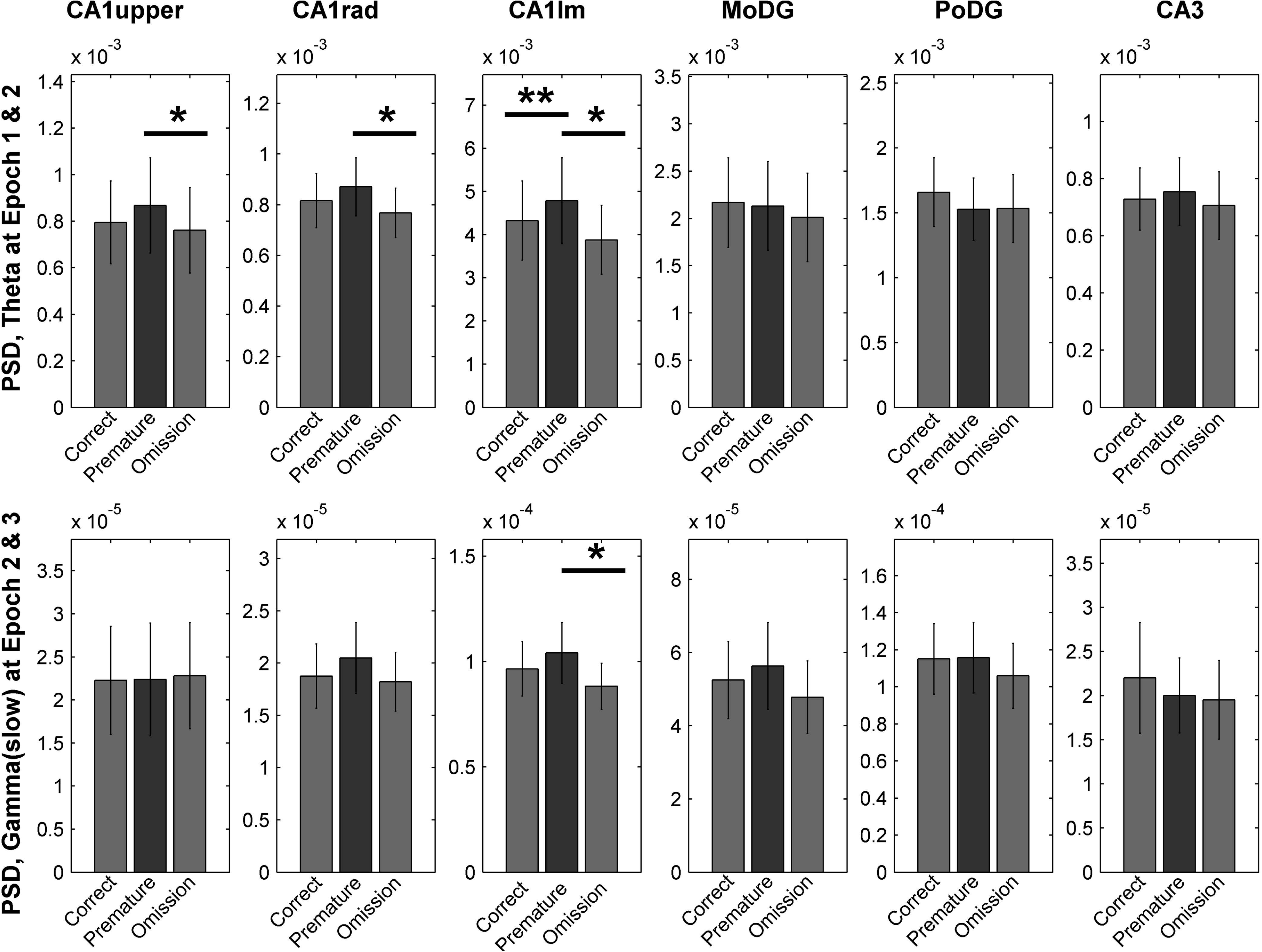

The TG-Phase alternations between the behavioral epochs 1, 2 and 3 were compared from data in which all the performance categories were pooled. The TG-Phase shift between the epochs was calculated, and the difference was tested with paired t-test. There was a significant TG-Phase shift in the slow-gamma (30–45 Hz) range and it was most prominent in the comparison between Epoch3 (after lever release) and Epoch1 (beginning of pressing) in the PoDG channel (Fig. 7 top-right). More precisely, the TG-Phase shifted later during Epoch3 compared to Epoch1 (PoDG TG-Phase 1–3 shift: confidence interval [23, 90]°, p = 0.002). A trend for a similar shift was observed between Epoch2 and Epoch 3 (PoDG TG-Phase 2–3 shift: confidence interval [-1, 55]°, p = 0.061). For the fast-gamma range (55–90 Hz) we also observed a significant shift in the CA1rad channel (Fig. 7 bottom-right). Interestingly, this TG-Phase shift had an opposite direction to that seen in the PoDG channel at slow-gamma range. The TG-Phase shifted earlier during Epoch3 compared to Epoch1 or 2 (CA1lower TG-Phase1–3 shift: confidence interval [-21, -76]°, p = 0.002; CA1lower TG-Phase12 shift: confidence interval [-14, -57]°, p = 0.004; Fig. 7 bottom-right column).

**Fig. 7.**
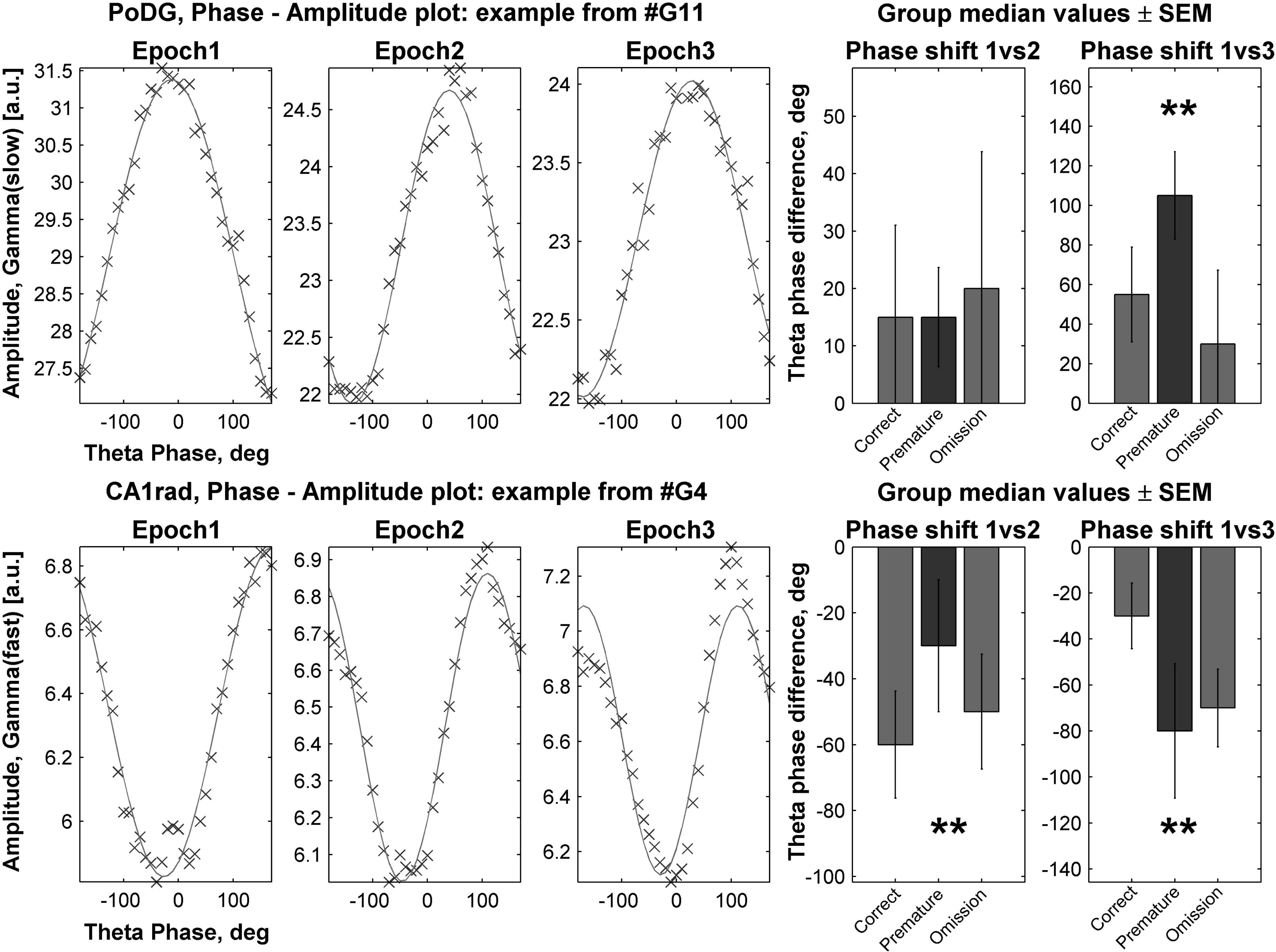

The task epoch and performance also affected the strength of theta-gamma modulation observed as changes in the modulation index (MI). First, the normalized modulation index (MInorm) was calculated by dividing the MI with the threshold (surrogate) value. Then, we computed ANOVA for repeated measures with two within-subject factors, epoch and performance. Interestingly, hippocampal locations which showed significant modulation of the MInorm by either task epoch or performance level were different from locations where effects on the TG-Phase shift were significant.

In the comparison of task epochs we observed in the slow-gamma range a reduction in the MInorm between Epoch1 and Epoch2, which was most prominent in the CA1upper channel (p = 0.009; Fig. 8 top left). In the CA1lm channel MInorm was significantly increased in Epoch3 compared to both Epoch 1 and 2 (Epoch 1 vs. 3, p = 0.034; Epoch 2 vs. 3 p = 0.032; Fig. 8 center and right). In contrast, in the fast-gamma range MInorm increased significantly between Epochs 2 and 3 in the CA1lm channel (Epoch 2 vs. 3, p = 0.013; Fig. 8 bottom left).

**Fig. 8.**
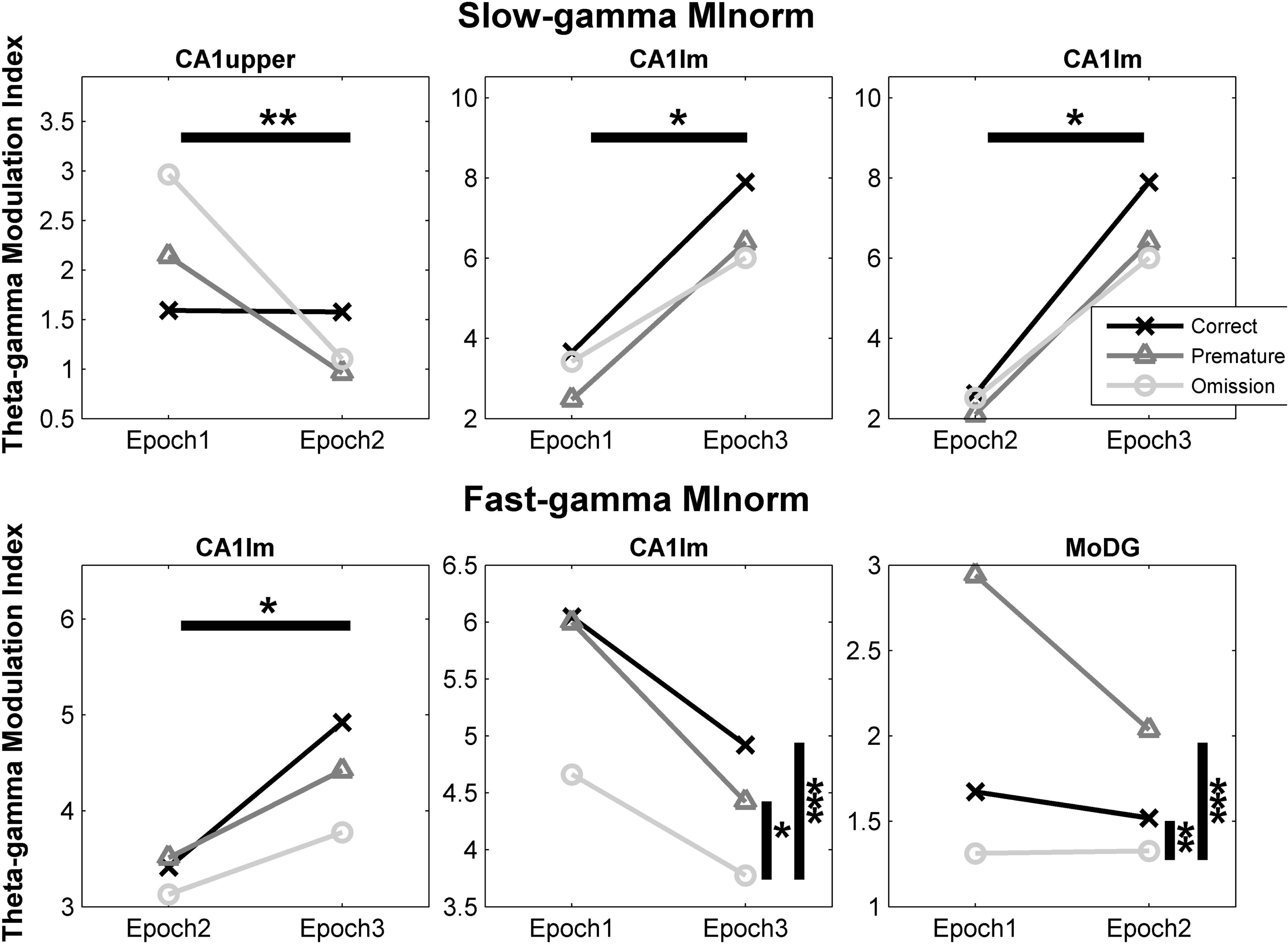

Interestingly, we also observed changes related to the task performance. Most notably, during omission trials the MInorm (for fast-gamma) in the CA1lm channel was smaller and reached significance when Epoch 1 vs. Epoch 3 were compared (LSD post-hoc test p = 0.001 [compared to correct trials] and p = 0.032 [compared to premature trials]; Fig. 8 bottom center). In addition, a significant performance effect for fast-gamma modulation strength was observed in the MoDG channel. The MInorm was higher during all error trials vs. correctly performed trials (Epoch 1 and 2, LSD post-hoc test p = 0.005 [compared to correct trials] and p = 0.009 [compared to > 4 s trials]; Fig. 8 bottom right).

## Discussion

The present experiment tested whether successful and or unsuccessful encoding of temporal requirements during lever pressing behavior could be predicted by observing activity of hippocampal theta or gamma rhythms. Indeed, we observed in addition to the behavioral changes in theta (6–10 Hz), slow-gamma (30–45 Hz) and fast-gamma (55–90 Hz) significant theta power differences between error and correct trials. Similarly, theta phase modulation of gamma amplitude in the two ranges (30–45 Hz & 55–90 Hz) (TG-Phase) as expressed as modulation index (MI) was specifically altered between different epochs but was also related to the performance accuracy.

We were able to confirm the previous findings on the relation between hippocampal theta and behavior. As expected, the strongest theta was observed after the lever pressing as the animal made the gross movements of releasing the lever and turning around, in line with the reported theta associated with voluntary movements (Vanderwolf, 1969; Whishaw and Vanderwolf, 1973; Buño, 1977). However, we detected hippocampal theta also during lever pressing, and that theta was stronger just before the lever release than in the beginning of lever pressing. Earlier studies have noted that theta in rats fluctuates before and after short**-**term lever pressing (Lopes da Silva and Kamp, 1969; Buño, 1977; Wyble et al., 2004), but it has remained disputable whether theta exists in immobile animals during prolonged lever pressing (Lopes da Silva and Kamp, 1969). Here we noticed that constant lever pressing does not mean immobility but involves small movements of the head, forelimb and sometimes the whole body. On the other hand, based on our observations immobility during lever pressing does not block hippocampal theta, which is consistent with findings of hippocampal theta in head-fixed rats (MacDonald et al., 2013). The increase in theta power during lever pressing could be linked to the sensorimotor model according to which hippocampal theta activity is crucial in continuously updated feedback of changing sensory (internal or external) conditions. Based on this model, hippocampal theta is implicated in initiation and maintain motor programs, and theta fluctuations before and after lever pressing relate to transition of motor programs (Bland and Oddie, 2001; Wyble et al., 2004).

Interestingly, theta power in the CA1lm channel showed correlation with the timing accuracy of the lever release. Specifically, theta power during lever pressing was higher than average during premature trials, but lower than average during omission trials. It is tempting to speculate that theta changes during ongoing lever pressing for several seconds imply that the hippocampus is trying to keep track on timing of the performance. However, we cannot exclude the possibility that more anxious behavior (with additional body movement) causes increase in theta power and a premature lever release. Although the exact mechanism of hippocampal theta generation is not clear, it is evident that the hippocampus may be able to generate theta intrinsically at least *in vitro* (Goutagny et al., 2009). Accordingly, current source density analyses in freely exploring rats have confirmed the co-existence of multiple theta current dipoles (Kamondi et al., 1998). Presumable sinks in lacunosum moleculare and stratum radiatum reflect excitatory inputs from CA3 area and entorhinal cortex while sources in stratum pyramidale reveal inhibitory inputs imposed by connections from the medial septum (Kocsis et al., 1999). Importantly, fibers originating from the medial septum terminate in essentially all fields of the hippocampal formation but are particularly prominent in the dentate gyrus (DG), and especially in the polymorphic layer. In contrast, fibers originating from the entorhinal cortex travel through the perforant path to terminate in all subdivisions of the hippocampal formation but mainly to the DG molecular layer (Witter and Amaral, 2004). Based on the known anatomical connectivity, we can conclude that observed changes in the PoDG most likely derive from changes in the medial septum input while theta increase in the CA1 area might also involve changes in the entorhinal input.

As for gamma, we observed that the gamma power was strongest in the beginning of the lever pressing and smallest just after lever release. This is particularly interesting because during lever pressing theta and gamma power developed into opposite directions. The task-related reduction of gamma power was most prominent in the PoDG channel which corresponds to the area with the strongest overall gamma power (Bragin et al., 1995). It has been proposed that there are two major gamma generators – one oscillator located in MoDG – PoDG layers of the dentate gyrus and a second in CA3 area. In addition, the DG oscillator is dependent on the input from the entorhinal cortex, while CA3 and CA1 gamma rhythms are coupled (Csicsvari, Jamieson et al. 2003). However, the functional significance of our findings cannot be directly derived from the known anatomical connectivity, since we still do not know exactly the conditions which enable hippocampal gamma to emerge, for instance compared to hippocampal theta (Leung et al., 1982; Buzsaki et al., 1983; Bragin et al., 1995).

However, gamma power is known to fluctuate between behavioral states, especially between REM sleep and waking. There is a clear increase in gamma power during REM sleep compared with active waking in the dentate gyrus. This change concurs with a decrease in CA1 gamma power while, interestingly, CA3 gamma power seems to remain the same (Montgomery et al., 2008). In addition, recent studies have implicated hippocampal gamma oscillations a role in more complex behavioral paradigms. In a spatial alternation task where the rat was required to sustain nose-poking for a 1-s fixation period, gamma-band oscillation in the CA1 area showed dynamical changes (Takahashi et al., 2014). In the beginning of fixation there was a strong fast-gamma (60–90 Hz) power followed by a reduction later during fixation, whereas the power of slow-gamma (30–45 Hz) increased later on in the fixation period. Interestingly, in error trials with premature fixation breaks, the changes in the power of slow- and fast-gamma were less evident during the shortest fixation periods (0 – 0.33 s) but the change in the gamma power of both frequency ranges was almost identical to correct trials in the longest premature breaks (0.67 – 1.0 s) (Takahashi et al., 2014). Similar changes in gamma power were also found in a hippocampal dependent odor-place association task (Igarashi et al., 2014). The power of the 20 – 40 Hz oscillation (including also beta frequency range) in the CA1 area showed dynamical changes during a 1-s fixation period during odor cue sampling. During this period the power of 20 – 40 Hz frequency band increased gradually during the cue sampling interval, peaked during the second half, and returned to baseline level as soon as the animal stopped sniffing (Igarashi et al., 2014). Slightly different results were obtained in a delayed nonmatch-to-place working memory task in a T-maze (Yamamoto et al., 2014). In the segments closest to the T-junction the power of fast gamma (65 – 140 Hz) was significantly greater than in other segments. However, such a difference was not seen in the slow-gamma range (25–50 Hz). In addition, this location-specific fluctuation in fast-gamma was dependent of the training phase, such that the fast-gamma power tended to decrease as the animals became over trained (Yamamoto et al., 2014).

Our finding of the gamma power reduction during lever pressing supports the earlier observation of reduced fast-gamma activity during fixation period but not the notion of simultaneous increase in slow-gamma activity (Takahashi et al., 2014). However, as shown the dynamics of gamma power fluctuations are complex and likely dependent on the task requirements (Igarashi et al., 2014). For instance, introducing sniffing during the fixation period seems to first enhance and later reduce not only the power of slow-gamma but also beta frequency (Igarashi et al., 2014). We also observed reduced gamma power during lever pressing with attention to a light signal but likely the constant lever pressing as such also produces more stable dynamical changes in gamma power than sniffing. Interestingly, the power of gamma is also dependent on the phase of the learning process, since overtraining seems to reduce observed fast-gamma (Yamamoto et al., 2014). Because we did not record brain activity during the learning phase we cannot conclude whether gamma dynamics changed during learning.

It is well established that the theta phase modulates gamma power (Buzsaki et al., 1983; Bragin et al., 1995). Therefore, observed gamma alternation during different task phase and performance in this study is not surprising. Further, we observed that theta**-**gamma modulation is not equally affected by the behavioral context in all hippocampal regions. Our results showed that the theta phase with the highest gamma power (TG-Phase) is shifted after lever release in PoDG and CA1lower channels, but these channels show a shift to opposite directions. Most likely this shift it is not related to gamma mechanisms *per se* but derives from a shift of theta current sinks and sources, and consequently, a shift in theta power, phase and coherence as has been earlier demonstrated in the context of a delayed spatial alternation task (Montgomery et al., 2009).

Theta-gamma modulation index (MI) has been shown to increase during learning and to predict the performance success in a memory task (Tort et al., 2009; Shirvalkar et al., 2010). However, as our present study shows, MI carries some additional information not related to theta or gamma power. Not all observed changes of MI in our study could be explained by theta or gamma power changes. First, MI is based on an adaptation of the Kullback–Leibler distance which has been shown to be insensitive to the absolute values of the amplitude envelope (Tort et al., 2010). Second, increase of MI (for both low- and fast-gamma) in CA1lm after lever release mirrored the increase of theta power in this layer. Although theta power increase was observed in other recording location as well, it was not associated with a significant MI change.

One interpretation of our findings is that MI changes especially in the CA1upper channel provide the context for attention and/or a temporal scale. This notion is supported by the observation that MI was attenuated during the lever pressing. However, the decrease of MI in the CA1lm channel and increase in the MoDG channel, implies location specific MI modulation.

